# Mechano-transduction *via* the pectin-FERONIA complex regulates ROP6 GTPase signaling in *Arabidopsis*

**DOI:** 10.1101/2021.03.17.435899

**Authors:** Wenxin Tang, Wenwei Lin, Binqi Li, Zhenbiao Yang

**Affiliations:** FAFU-UCR Joint Center for Horticultural Biology and Metabolomics Center, Haixia Institute of Science and Technology, Fujian Agriculture and Forestry University, Fuzhou, Fujian, China; Institute of Integrative Genome Biology and Department of Botany and Plant Sciences, University of California, Riverside, CA 92521

## Abstract

During growth and morphogenesis, plant cells respond to mechanical stresses resulting from spatiotemporal changes in the cell wall that bear high internal turgor pressure. Microtubule (MT) arrays are re-organized to align in the direction of maximal tensile stress to guide the synthesis of cellulose, reinforcing the local cell wall. However, how mechanical forces regulate MT re-organization remains largely unknown. Here, we demonstrate that mechanical signaling that is based on the CrRLK1L subfamily receptor kinase FERONIA (FER) regulates the reorganization of cortical MT in cotyledon epidermal pavement cells (PC) in *Arabidopsis*. Recessive mutations in *FER* compromised MT response to mechanical perturbations such as single cell ablation, compression and Isoxaben treatment in these pavement cells. These perturbations promoted the activation of ROP6 GTPase that acts directly downstream of FER. Furthermore, defects in the ROP6 signaling pathway negated the reorganization of cortical MTs induced by these stresses. Finally, reduction in highly demethylesterified pectin, which binds the extracellular malectin domain of FER and is required for FER-mediated ROP6 activation, also impacted mechanical induction of cortical MT reorganization. Taken together our results suggest that the FER-pectin complex senses and/or transduce mechanical forces to regulate MT organization through activating the ROP6 signaling pathway in *Arabidopsis*.

## INTRODUCTION

Cellular organisms experience external forces as well as internal mechanical cues, which need to be integrated with chemical signals to coordinate cell behaviors in space and over time. Plant cells are under tremendous mechanical stresses from strong internal turgor pressure. As plant cells are attached to each other via the cell wall, supracellular patterns of physical forces are generated during growth, causing large tension in the epidermis. The resulting forces may provide a coordinating supracellular signal to regulate cell division and cell growth/expansion and plant morphogenesis from the cell to organ level [1–4]. Changes in growth pattern and cell expansion requires corresponding local or global modification of the cell wall, resulting in spatiotemporal changes in mechanical stresses from turgor pressures. Plant cells must monitor these mechanical changes to assure the cell wall homeostasis and integrity and to coordinate with cell growth and morphogenesis [5, 6]. The MSL family members have been shown to monitor osmotic shocks for the maintaining of cell viability [7, 8], while CrRLK1L members might be candidates for the sensors that monitor mechanical forces for cell wall modification in plants [5, 9]. However, direct evidence for their mechanosensing capacity is lacking, and thus the molecular mechanisms for mechanical sensing and signaling in plants remain poorly understood.

As in animal cells, the cytoskeleton plays an important role in mechanical responses in plant cells. Cortical microtubules (MT) were found to align with maximal stress in plants [10, 11]. Increasing evidence supports a role for mechanical forces in the regulation of MT orientation in plant cells [12–17]. MT in turn direct the synthesis of cellulose to reinforce the cell wall in countering the mechanical stress. Pavement cells (PC) in the cotyledon and leaf epidermis provide an attractive system for studying MT-based mechanical feedback regulation of cell morphogenesis [18]. In most angiosperms, leaf epidermal PC exhibit jigsaw puzzle shapes with interdigitated indention and lobe-like outgrowths in the anticlinal direction. Two distinct, but not necessarily exclusive, models have been proposed to initiate the interdigitated cell shape pattern. Auxin is thought to be the chemical signal, as it is important for and promotes the lobing of PCs by activating ROP/Rac GTPase-dependent local cytoskeletal reorganization [19–24]. In particular, cortical MTs enriched in PC indentation regions guide localized deposition of stiff cellulose microfibrils (CMF), thus inhibiting outgrowth [20, 22, 25–28], and this localized MT organization is promoted by katanin-based MT severing promoted by ROP6, which is activated by cell surface auxin sensing [22, 24, 29].

In parallel or consequently, mechanical stress may trigger a MT-based feedback mechanism for the cellulose-based cell wall reinforcement, and thus the maintenance of pavement cell shape [18, 30]. This stress-induced positive feedback loop may be initiated by auxin/ROP pathway described above. In an alternative non-exclusive scenario, mechanical and chemical polarities may emerge early on in the anticlinal walls of PC, before they start waving [31]. Mechanical forces were recently found to modify MT organization in leaf PC [32]. It has been proposed that stress in the outer wall may amplify such initial deformations [31, 33], thereby maintaining mechanical stress at a relatively low level, while increasing cell volume [34]. Identification of the unknown molecule that senses the mechanical stress would help to understand how the mechanical signal activates MT reorganization and whether it acts independently from or coordinately with the auxin signal.

FER (Feronia), CrRLK1Ls family (Catharanthus roseus RLK1-like kinase family) [35], regulates multiple processes during plant growth, such as plant immunity [36–38], cell elongation [39, 40], root and root hair growth [41, 42], cell morphogenesis [43] and seed development [44]. Most of these processes are linked to cell wall homeostasis and/or cell wall integrity (CWI) maintenance. Moreover, FER is required for mechanically induced changes in Ca^2+^ [4]. CrRLK1L members contain two extracellular carbohydrate-binding malectin (Mal) domains [35]. Feng et al recently showed that FER binds pectin *in vitro* through its extracellular domain and maintains cell-wall integrity during salt stress [45]. Furthermore, Lin et al showed that FER senses highly demethyesterified pectin to directly activate the ROP6 signaling pathway during PC morphogenesis in *Arabidopsis* [46] (accompanying manuscript). Here we show that FER participates in the sensing and/or signaling of mechanical stress that activate the ROP6 signaling pathway to induce MT rearrangement in *Arabidopsis* cotyledon epidermis.

## RESULTS

### The *fer-4* Mutation Compromises Sensitivity to Mechanical Stress-induced MT Rearrangement

Recessive *fer* alleles exhibit severe pavement cell (PC) shape defects, such as wider indentation, less shallow lobes, indicative of weaker cell wall in the indentation regions of these cells [43, 46]. Given demethylesterified pectin’s binding to and requirement for the activation of FER [46], the cell wall reinforcement may be mediated by FER’s sensing of mechanical stress via its interaction with pectin. In animals, extracellular matrix (ECM)-integrin (ECM receptor) plays an important role in sensing mechanical forces [47]. Binding of the cell wall to Wsc1 is also proposed to be essential for its sensing of mechanical stress in yeast [48–51]. Here we tested whether FER is involved in sensing the mechanical stress using a MT re-organization assay, as the reorganization of cortical MT (CMT) has been used as a readout for mechanical stress [17, 30]. Cortical MT organization into ordered arrays is indicated by MT anisotropy measured by FibrilTool, an ImageJ plug-in to quantify fibrillar structures in raw microscopy images [52]. MT has often been labeled with fluorescence protein tagged-tubulin, but we found GFP-tubulin’s labeling of cortical MTs in *fer-4* PCs was quite weak, making the measurement of MT anisotropy quite difficult. Thus we used GFP-MAP4, which gives clear labeling of MTs both in wild type and *fer* mutants. As shown previously with GFP-tubulin labeled MT, mechanical stress induced a dramatic increase in the anisotropy of MT labeled with GFP-MAP4. MT anisotropy in *fer-4* PCs (cultured in ½ MS with 1% agar) was reduced to nearly 50% (from 0.058±0.013 to 0.034±0.010) in the indentation region when compared to wild type PC, consistent with wider indentation regions in *fer-4* PC (Figure S1) [46].

To assess whether FER regulates microtubule response to mechanical stress, external mechanical forces generated by ablation, compression and Isoxaben treatments were applied on cotyledons to alter stress patterns [30]. Firstly, we carried out single cell ablation by high-dosage UV light radiation on the epidermis of cotyledons from seedlings cultured in ½ MS with 1.0% agar. Maximal stress directions are predicted to become circumferential to the ablation site independent of cell geometry [30]. In WT, 4-8 hours after ablation, the orientation of cortical MT arrays in PCs surrounding the ablation site changed to a circumferential rearrangement (Figure 1A), as reported previously [30]. The anisotropy of cortical MT was greatly increased (from 0.07±0.038 to 0.16±0.061 and 0.18±0.060 at 4 and 8 hrs after ablation, respectively). However, when a single PC was ablated in *fer-4* cotyledons, the anisotropy was only slightly increased from 0.038±0.019 to 0.046 ±0.023 and 0.058±0.024 at 4 and 8 hrs, respectively (Figures 1A, 1B and 1E). Thus, ablation induced MT rearrangement was essentially lost in the *fer-4* mutant.

**Figure 1.**
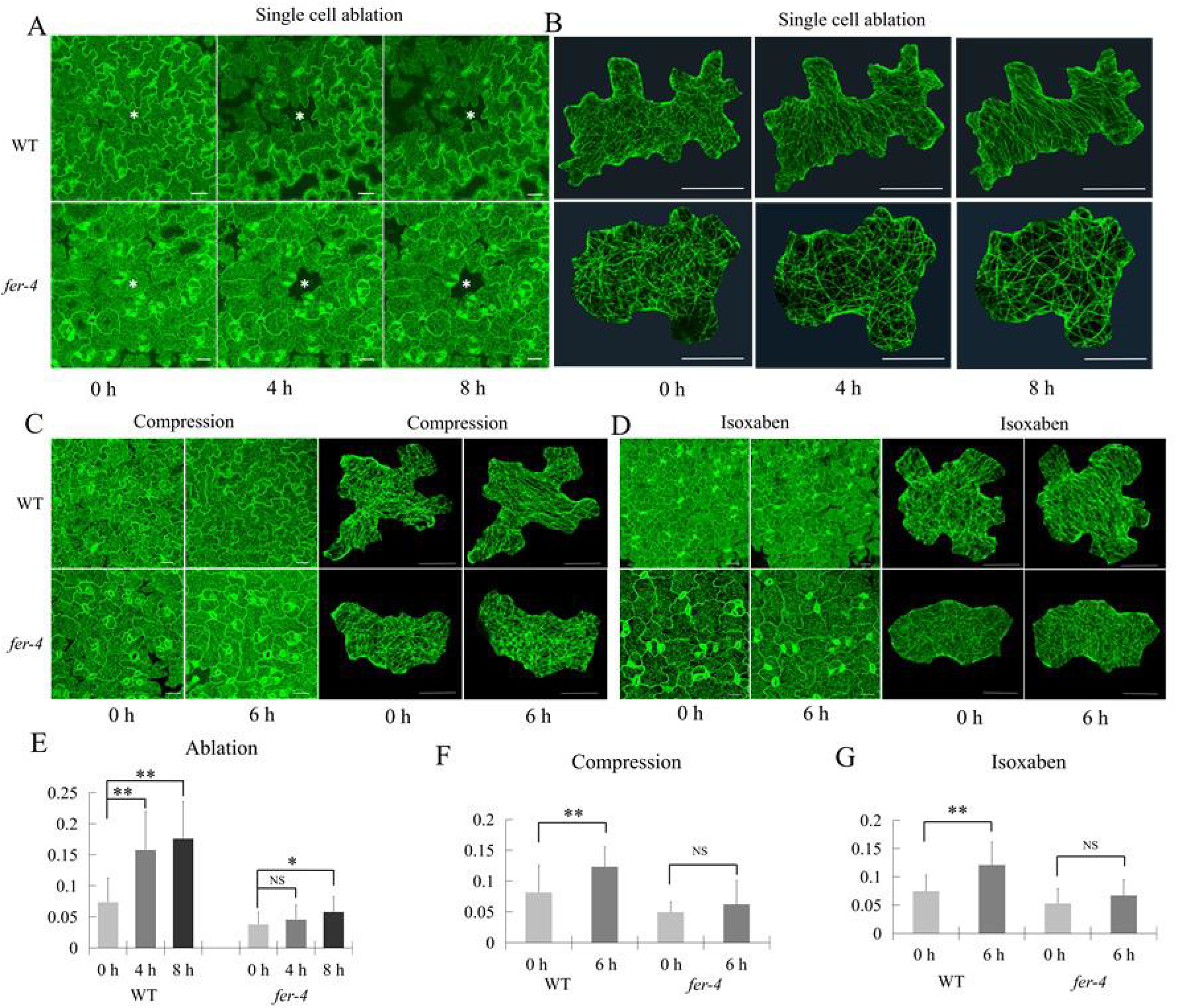
*fer-4* PCs exhibit reduced sensitivity to mechanical stresses in inducing MT rearrangement. Cotyledons from Col-0 (WT) and *fer-4* expressing GFP-MAP4 were subject to different mechanical stresses induced by single cell ablation (A and B), compression (C) and isoxaben treatment (D) and cortical MT anisotropy were measured (E, F, and G, respectively). (A) MT rearrangement after single cell ablation treatments on 3 DAG (days after germination) cotyledon PCs from WT (GFP-MAP4) and *fer-4* (*fer-4 x GFP-MAP4*). In WT, ablation of single PC resulted in the circumferential distribution of MT arrays around the site of physical perturbations (White asterisk marks site of ablation). In *fer-4* mutant, MT rearrangement induced by single cell ablation nearly lost. Scale bar 50 μm. (B) Representative PCs from A were enlarged to show clear MT arrangements. Scale bar, 50 μm. (C) MT rearrangement after 6 hrs compression treatment on WT and *fer-4* 3 DAG cotyledon PCs. Representative individual cells are shown in the right panels. Scale bar, 50 μm. (D) MT rearrangement after 6 hrs Isoxaben (20 μM) treatments on WT and *fer-4* 3 DAG cotyledon PCs. Representative individual cells are shown in the right panels. Scale bar, 50 μm. (E) Quantification of MT Anisotropy in A. Two layers of PCs surrounding the ablated cell were used for quantification. Data are mean degrees from >20 independent cells ±SE. Asterisks indicate the significant difference (*p<0.05, **p<0.01, NS, no significant difference, Student’s *t*-test) between the wild type and the mutants in the above assays. (F-G) Quantification of MT Anisotropy in C and D. PCs from the same region of cotyledons (just below the tip) were quantified. Data are mean degrees from >20 independent cells ±SE. Asterisks indicate the significant difference (*p<0.05, **p<0.01, NS, no significant difference, Student’s *t*-test) between the wild type and the mutants in the above assays.

Application of compressive forces resulted in an increase in overall stress and hyper-alignment of MT [30]. We directly applied compressive forces to PCs by using a coverslip pressed on the surface of the cotyledons and kept in place using adhesive silicone applied on the margins [32]. The rearrangement of cortical MT became more aligned by 6 hrs in PCs from WT cotyledons but not from *fer-4* (Figures 1C and 1F). Isoxaben, which inhibits cellulose synthesis and thus increases tensile stress on the cell wall, was shown to induce hyper-bundling and hyper-alignment of cortical MT along the predicted directions of maximal stress in shoot meristem cells [53]. Treatment of 3-day-old seedlings with 40 μM isoxaben for 6 hrs led to a sizeable increase in the anisotropy of MT (Figures 1D and 1F). In *fer-4* mutant, MT kept randomly aligned 6 hrs after the treatment (Figures 1D and 1F).

### Mechanical stresses promote ROP6 GTPase activation

Katanin-mediated MT severing is required for MT responses to mechanical stresses in the shoot apical meristem and leaf PC in *Arabidopsis* [30, 54]. In *Arabidopsis* PCs, cortical MT organization is regulated by the ROP6 signaling pathway acting upstream of katanin [22, 29], and FER activates ROP6 directly via their interactions with RopGEF14 [46]. Thus, we hypothesized that mechanical stress activates the ROP6 pathway *via* FER to promote the katanin-mediated MT rearrangement. To test this hypothesis, we first determined whether ROP6 activation is mediated by mechanical stress. We treated 7 DAG (days after germination) GFP-ROP6 seedlings [24] with 40 μM Isoxaben, which induces an external mechanical stress overriding the internal developmental stress pattern. The amount of active GFP-ROP6 protein increased by 50-100% in 1 h to 2 h after Isoxaben treatments compared with untreated control (0 h, Figure 2A). This effect is specific for ROP6, because the amount of active GFP-ROP2 protein was not significantly affected by the same treatment in GFP-ROP2 transgenic seedlings (Figure 2B). Different concentrations of Isoxaben will generate different strengths of mechanical forces. We found that 10-40 μM Isoxaben were the optimal concentrations for the induction of ROP6 activity (Figure S3A). We confirmed that these concentrations of Isoxaben induced cell shape changes (Figure S2).

**Figure 2.**
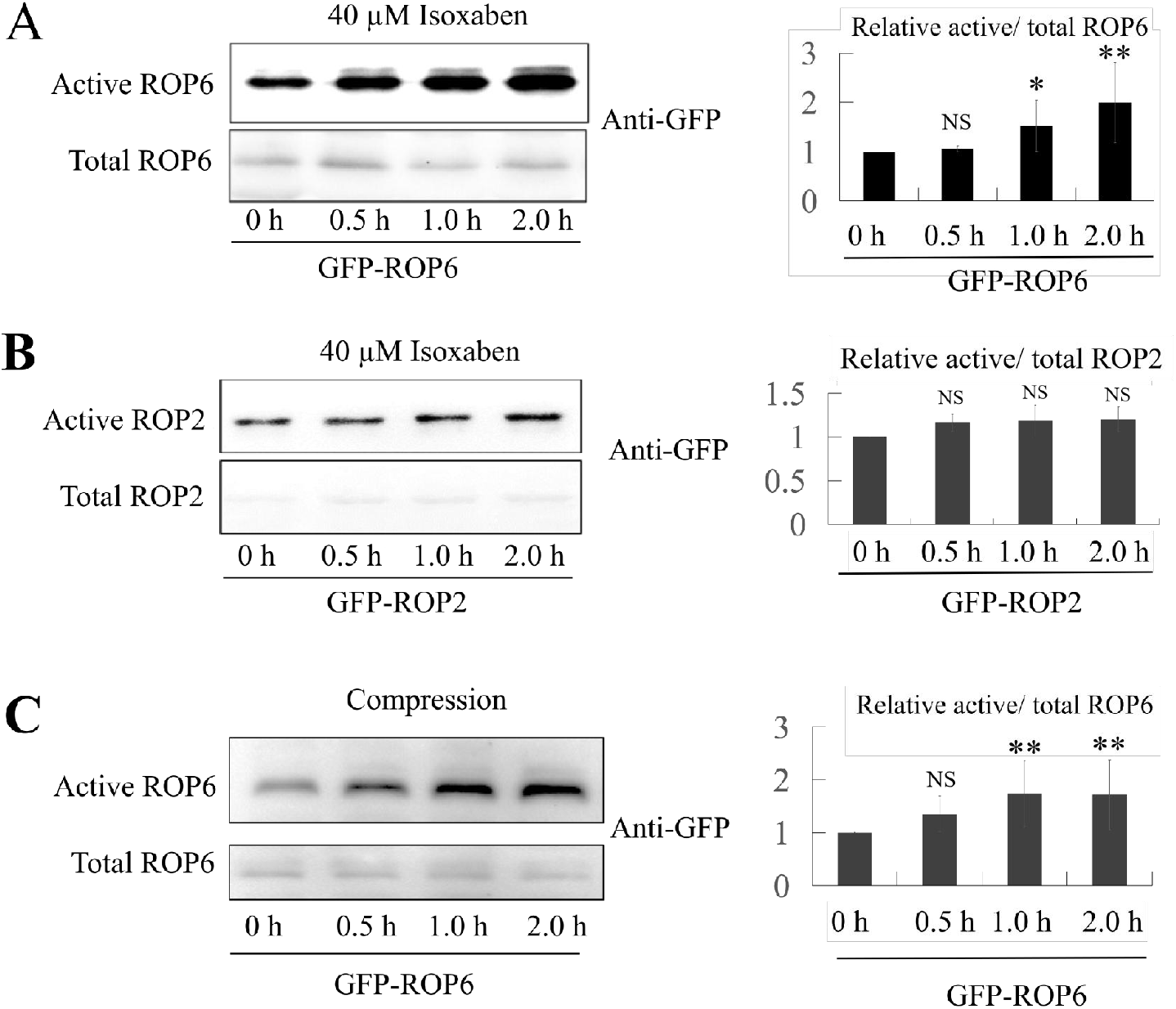
Mechanical stresses induce the activation of ROP6 but not ROP2. (A) Western blots based ROP6 activity measurement in 7 DAG seedlings of GFP-ROP6 with isoxaben treatments in different times. ROP6 activation by Isoxaben was analyzed by pull-down assay, as described previously [27]. Quantification of relative active GFP-ROP6 level (amount of GTP-bound GFP-ROP6 divided by amount of total GFP-ROP6) to control (as “1”) is shown. Data are mean activity levels from three independent experiments ±SD. *p<0.05, **p<0.01, NS, no significant difference, Student’s *t*-test. (B) ROP2 activity in GFP-ROP2 with and without isoxaben treatments in different times. ROP2 activity can be slightly induced by 40 μM Isoxaben. Data are mean activity levels from three independent experiments ±SD. NS, no significant difference, Student’s *t*-test. (C) ROP6 is activated by compression. ROP6 activity in 3 weeks leaves of GFP-ROP6 with compression treatment in different times. Data are mean activity levels from three independent experiments ±SD. **p<0.01, NS, no significant difference, Student’s *t*-test.

To directly test whether ROP6 activity is induced by mechanical stress, we next assessed the effect of externally applied compression on ROP6 activity. Compression was applied on 3-weeks old leaves. Four glass slides were placed on one half of a leaf as compression treatment, and the other half was free of external pressure as control. At various times, the glass slides were removed, and the leaf was dissected into two halves and immediately frozen in liquid nitrogen for ROP6 activity assay. The compression induced an increasing ROP6 activation over the times of treatment (Figure 2C). After 2 hours of compression, the amount of active GFP-ROP6 increased to about 2 times compared with the untreated control (0 h, Figure 2C). When compression was applied on Col-0 leaves, the activation of native ROP6 also detected after 2 hours of compression (Figures S3B and S3C). These results clearly demonstrate that mechanical stresses promote ROP6 activation.

### FER and RopGEF14 Are Required for the ROP6 Activation Induced by Mechanical Stress

We next determined whether FER is required for the activation of ROP6 induced by mechanical stress. GFP-ROP6 and *fer-4x GFP-ROP6* lines were used for Isoxaben and compression treatments as described above. In contrast to a 1-fold increase in the amount of active GFP-ROP6 after 7 DAG GFP-ROP6 seedlings were treated with Isoxaben for 1-2 hrs, no significant increase in GFP-ROP6 activity was observed in *fer-4xROP6-GFP* seedlings with the same treatments (Figure 3A). Similar results were obtained when ROP6 activity was assayed in Col-0 and *fer-4* lines using anti-ROP6 to detect ROP6 (Figure S4A).

**Figure 3.**
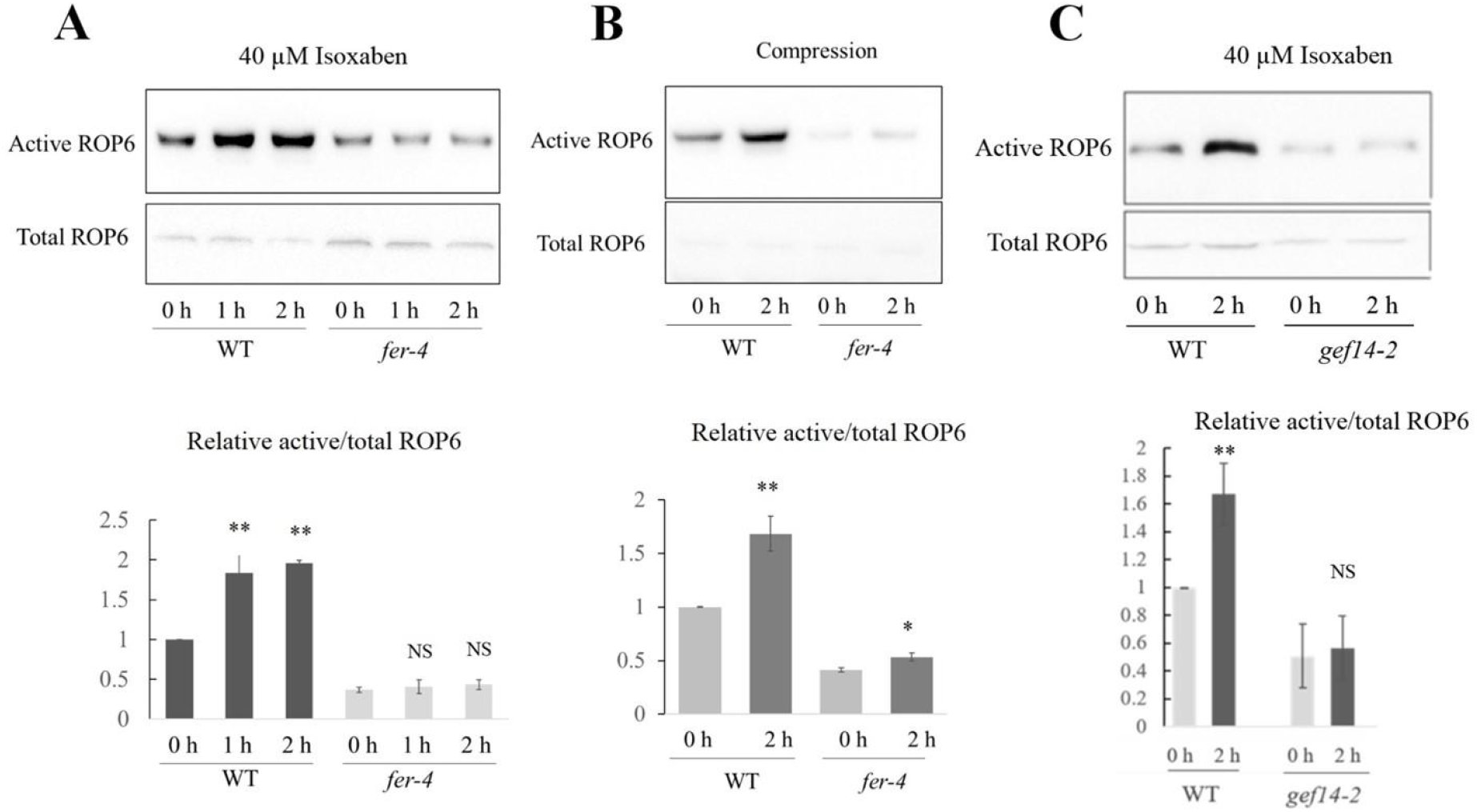
FER is required for mechanical stress on ROP6 activation. (A) ROP6 activity in 7 DAG seedlings of GFP-ROP6 and *fer-4 x GFP-ROP6* with and without isoxaben treatments for different times. ROP6 activity can be induced by 40 μM Isoxaben in GFP-ROP6 but not in *fer-4 x GFP-ROP6*. Data are mean activity levels from three independent experiments ±SD. **, P<0.01; NS, no significance (B) Compression induced ROP6 activation is lost in *fer-4*. Compression was applied on leaves from 3 weeks old control (GFP-ROP6) and *fer-4* (*fer-4 x GFP-ROP6*) plants. After 2 hrs of compression, leaves were loaded for ROP activity measurement. Data are mean activity levels from three independent experiments ±SD. *, p< 0.05; **, P<0.01 (C) ROPGEF14 is required for Isoxaben induced ROP6 activation. ROP6 activity in 7 DAG seedlings of WT and *gef14-2* with 40 μM isoxaben treatments for 2 hours. ROP6 activity can be induced by 40 μM Isoxaben in WT but not in *gef14-2*. Data are mean activity levels from three independent experiments ±SD. **, P<0.01; NS, no significance

We also investigated whether FER is required for the ROP6 activation induced by compression in 3 weeks old GFP-ROP6 and *fer-4xGFP-ROP6* leaves. As shown in Figure 3, 2 hrs of compression induced GFP-ROP6 activation by two folds compared with mock treatment in GFP-ROP6 leaves. However, in *fer-4xGFP-ROP6* leaves, the amount of active ROP6 decreased in the control, and compression failed to induce ROP6 activation (Figure 3B).

FER activates ROP6 signaling through direct interaction with the ROP6 activator ROPGEF14 in the cotyledon epidermis [46]. Therefore, ROPGEF14 is expected to be required for mechanical stress induced ROP6 activation. To confirm this, Isoxaben and compression were applied on the 7-day old seedlings and 3 weeks-old leaves from the *gef14-2xGFP-ROP6* and GFP-ROP6 lines [46], respectively. As expected, the *ropgef14-2* knockout mutation greatly compromised GFP-ROP6 activation induced by both Isoxaben and compression (Figures 3C and S4B). Taken together, we show that the FER-ROPGEF14 complex is required for the induction of ROP6 activation by mechanical forces.

### ROP6 Signaling Is Required for the MT Rearrangement Induced by Mechanical Stress

Because FER-dependent mechano-transduction activates MT rearrangement and ROP6 signaling, which was previously shown to promote the organization of cortical MT in PCs [22], it is reasonable to hypothesize that the FER-based mechano-transduction regulates MT rearrangement *via* ROP6 signaling. To test this hypothesis, we investigated whether ROP6 signaling is required for mechanical stress induced MT rearrangement. We previously showed that RopGEF14 and RIC1 act as ROP6 activator and effector to regulate MT organization, respectively [22, 46]. Thus, three mutants *gef14-2, rop6-1* and *ric1-1* were used to assess the role of ROP6 signaling in the mechanical induction of MT rearrangement. Unlike *fer-4*, all these three mutants only exhibit relatively mild defects in PC morphogenesis, likely due to the existence of another ROP or signaling pathway that is functionally overlapping with ROP6 in the promotion of MT ordering during PC morphogenesis. Thus, we expected these mutants to be less compromised in the response to external forces than *fer-4*. Indeed 3 DAG cotyledons from *gef14-2, rop6-1* and *ric1-1* mutants all appeared to have normal sensitivity to ablation-induced MT rearrangement when seedlings were cultured in ½ MS medium with 1% agar (Figure S5). We reasoned that lowering tensile stress would reduce the response of MT rearrangement to mechanical stress and thus condition the *gef14-2*, *rop6-1* and *ric1-1* mutants to be less sensitive to the weaker mechanical stress. Older cotyledons, which have thicker and stronger cell wall, are expected to experience less internal tensile stress. In addition, cell wall tension is reduced when seedlings are cultured in higher concentrations of agar (2.5%) than in regular agar concentration (1%) [17]. Therefore, we cultured *gef14-2*, *rop6-1* and *ric1-1* mutants to 7 DAG in both regular and high concentrations of agar (1% and 2.5%). We found that the *gef14-2* mutation greatly reduced the sensitivity to single cell ablation in inducing MT rearrangement in 7 DAG cotyledons under normal (1%) agar conditions (Figures 4A and 4C). Both *rop6-1* and *ric1-1* mutants exhibited less response to ablation when cultured in ½MS with 2.5 % agar (Figures 4 B and 4D). To further confirm the importance of the ROP6 signaling pathway in mechanical stress-induced MT rearrangement, we incubated 7 DAG *gef14-2*, *rop6-1* and *ric1-1* in high concentration of agar (2.5%) in ½ MS with 40 μM Isoxaben. MT rearrangement were induced and the anisotropy value ratio (4h/0h) was more than 2 in WT, whereas in *gef14-2*, *rop6-1* and *ric1-1* the anisotropy value ratios (4h/0h) were less than 1.5 (Figures 4E and 4F). Taken together, the ROP6 pathway is required for MT organization response to mechanical stress.

**Figure 4.**
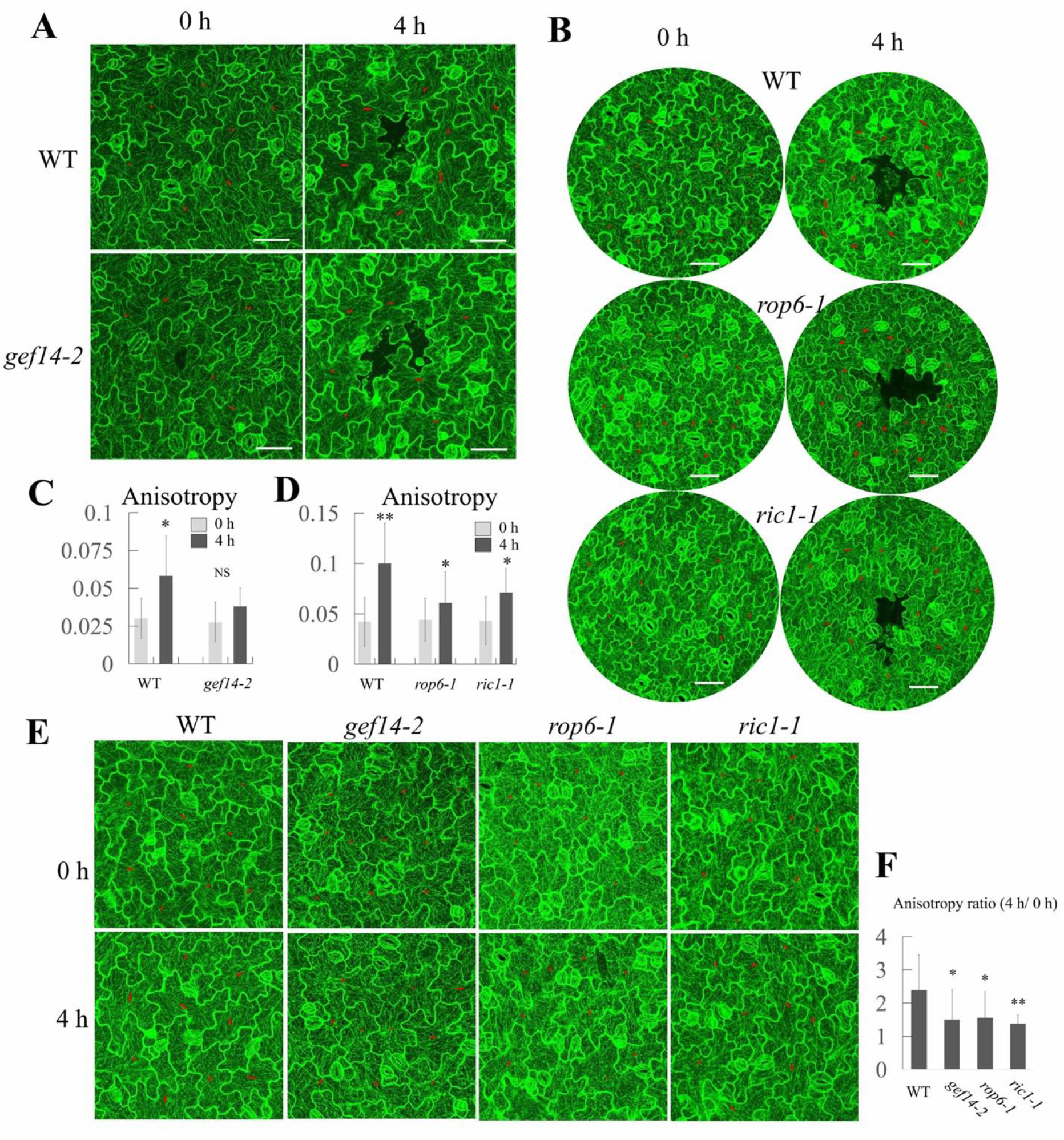
ROP6 signaling is required for mechanical stress induced MT rearrangement. (A) MT rearrangements after single cell ablation in WT and *gef14-2* PCs. 7 DAG cotyledons grown in ½ MS with normal concentration of agar (1.0%) were observed by confocal microscopy. Red line represents the anisotropy of MT. Two layers of PCs around ablation point were quantified. Bar=50 μm. (B) MT rearrangements after ablation in WT, *rop6-1* and *ric1-1* PCs. 7 DAG cotyledons grown in ½ MS with high concentration of agar (2.5%) were observed in confocal microscopy. Red line represents the anisotropy of MT. Two layers of PCs around ablation point were quantified. Bar=50 μm. (C) Anisotropy values of MT in (A). *gef14-2* was less sensitive to ablation-induced anisotropy increase. Data are mean degrees from >50 independent cells ±SE. *, p<0.05; NS, no significance (D) Anisotropy value of MT in (B). Both *rop6-1* and *ric1-1* were less sensitive to ablation-induced anisotropy changes compared to WT. Data are mean degrees from >50 independent cells ±SE. *, p<0.05; **, p<0.01. (E) MT rearrangements after 40 μM Isoxaben treatment in WT, *gef14-2*, *rop6-1* and *ric1-1* PCs. 7 DAG cotyledons grown in ½ MS with high concentration of agar (2.5%) were observed by confocal microscopy. Red line represents the anisotropy of MT. PCs at the same position of cotyledon were quantified. Bar, 50 μm. (F) Anisotropy value ratios (4 h / 0 h) of MT in E. *gef14-2*, *rop6-1* and *ric1-1* were less sensitive to 40 μM Isoxaben treatment in the induction of anisotropy change compared to WT. Data are mean degrees from >50 independent cells ±SE. B.*, p<0.05; **, p<0.01

### Highly Demethylesterified Pectin Is Required for ROP6 Signaling Induced by Mechanical Stress

The connection between a transmembrane receptor and extracellular matrix provides a common mechanism for sensing mechanical forces in animal and fungal cells [55]. FER binds highly demethylesterified pectin through its extracellular MALA domains to ROP6 signaling in during PC morphogenesis [46]. Indentation regions of the anticlinal wall in PC are enriched in highly demethylesterified pectin [46] and undergo highest mechanical stress [30]. Thus, it is reasonable to propose that the FER-pectin connection senses the mechanical stress in these indentation regions, generating a positive feedback regulation of ROP6 signaling and MT rearrangement and subsequent microfibril deposition for the re-enforcement of the indentation regions. To test this hypothesis, we investigated whether highly demethylesterified pectin is required for mechanical force sensing. To assess the impact of pectin demethylesterification levels, we used a transgenic line overexpressing pectin methylesterase inhibitor 1 (PMEI1 OE), which contains reduced pectin demethylesterification [46]. First, we performed single cell ablation on 3 DAG cotyledon epidermis of the PMEI1 OE line. Four hrs after ablation, cortical MT of PCs became greatly rearranged in WT (Figure 5A). However, compared to WT, PMEI1OE PCs showed less response to ablation (Figure 5A). In WT, the anisotropy was increased nearly by 2 folds from 0.041±0.007 (0 hr) to 0.116±0.026 at 4 hrs after ablation. In contrast, in PMEI1OE the anisotropy was increased by one fold from 0.031±0.006 (0 h) to 0.066±0.011 (4 h; Figure 5B).

**Figure 5.**
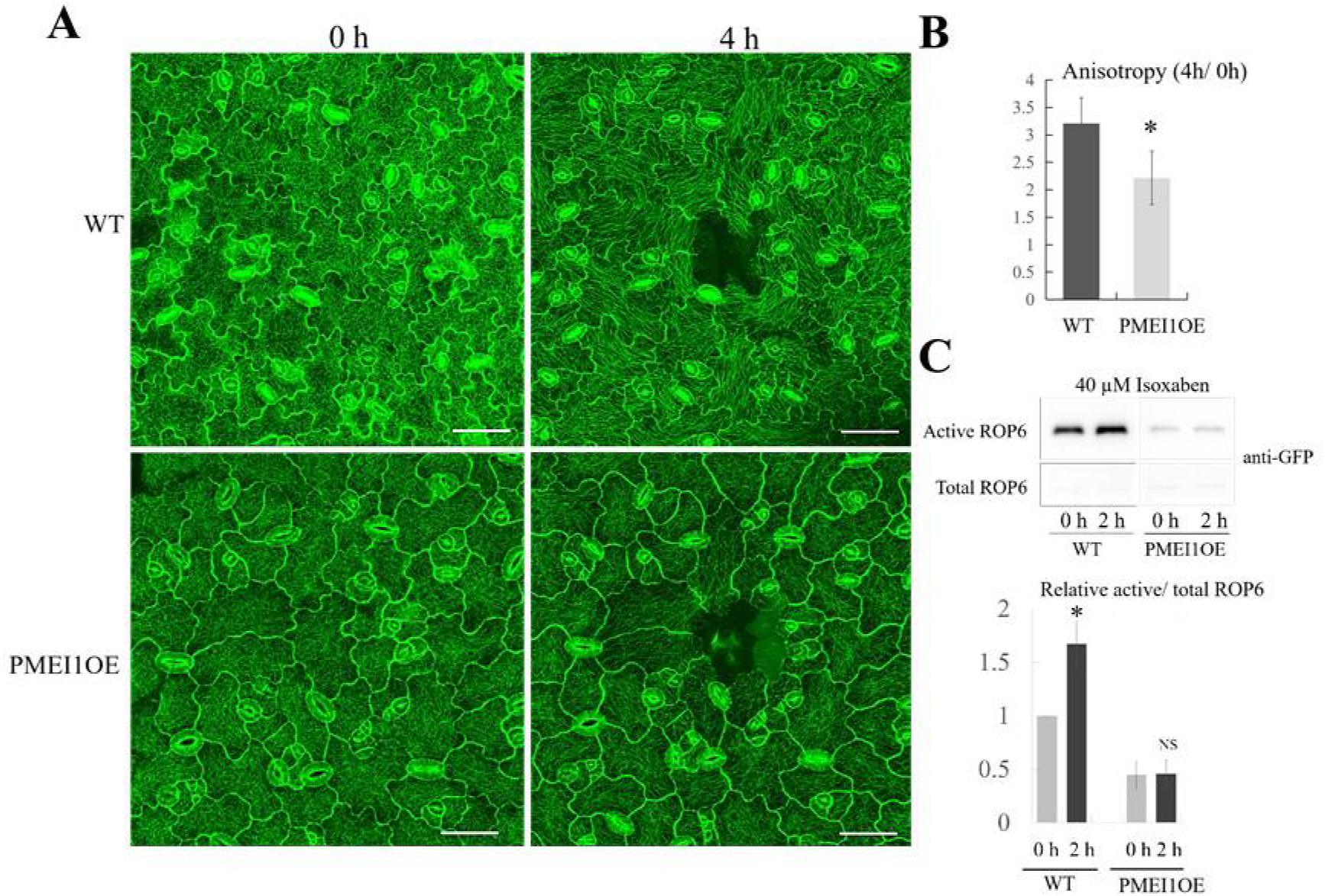
Demethylesterified pectin is important for the sensing of mechanical stress to induce MT rearrangement and ROP6 activation. (A) CMT rearrangement after single cell ablation on 3 DAG cotyledons from the GFP-MAP4 and PMEIOE/GFP-MAP4 lines. Scale bar=50 μm (B) Quantification of MT Anisotropy in A. Two lays of PCs surrounding the ablated cell were chosen for quantification. Scale bar=50 μm. Data are mean degrees from >20 independent cells ±SE. Asterisks indicate the significant difference (*p<0.05, Student’s *t*-test) between the wild type and the mutants in above assays. (C) ROP6 activity in 7 DAG seedlings of WT and PMEI1OE with and without 40 μM Isoxaben treatment. Data are mean activity levels from three independent experiments ±SD. *, P<0.05; NS, no significance

Next we investigated whether pectin demethylesterification affects ROP6 activation induced by mechanical stress. ROP6 activity in seedlings of WT and PMEI1OE with and without 40 μM Isoxaben treatment was assayed. As expected, Isoxaben induced ROP6 activity was greatly reduced in PMEI1OE line (Figure 5C). These results suggest that highly demethylesterified pectin is required for ROP6 signaling induced by mechanical stress.

## DISCUSSION

Our current study together with the accompanying paper [46] provides strong support for the hypothesis that FER participates in sensing and/or transducing mechanical signal to directly regulate MT reorganization in the control of cell expansion and cell shape formation. First, we showed that both FER and its cell wall binding partner highly demethylesterified pectin are both required for MT re-orientation induced by mechanical stress. Second mechanical stress activates ROP6 signaling and this activation requires both FER and highly demethylesterified pectin. Third, the ROP6 signaling components, including ROP6 itself, its activator RopGEF14 that directly interacts with FER, and its effector RIC1 that directly regulate MT organization, are all important for the MT re-arrangement induced by mechanical stress (Figure 6) [46]. To our knowledge, the pectin/FER-ROP-MT pathway is the most complete mechano-transducing pathway connecting mechanical stress to the cellular responses described in plants to date. Nonetheless, the confirmation of FER’s role as a direct mechanosensor requires single molecule mechanosensing assays in which mechanosensing activity can be measured when forces are applied to the single FER molecule.

**Figure 6.**
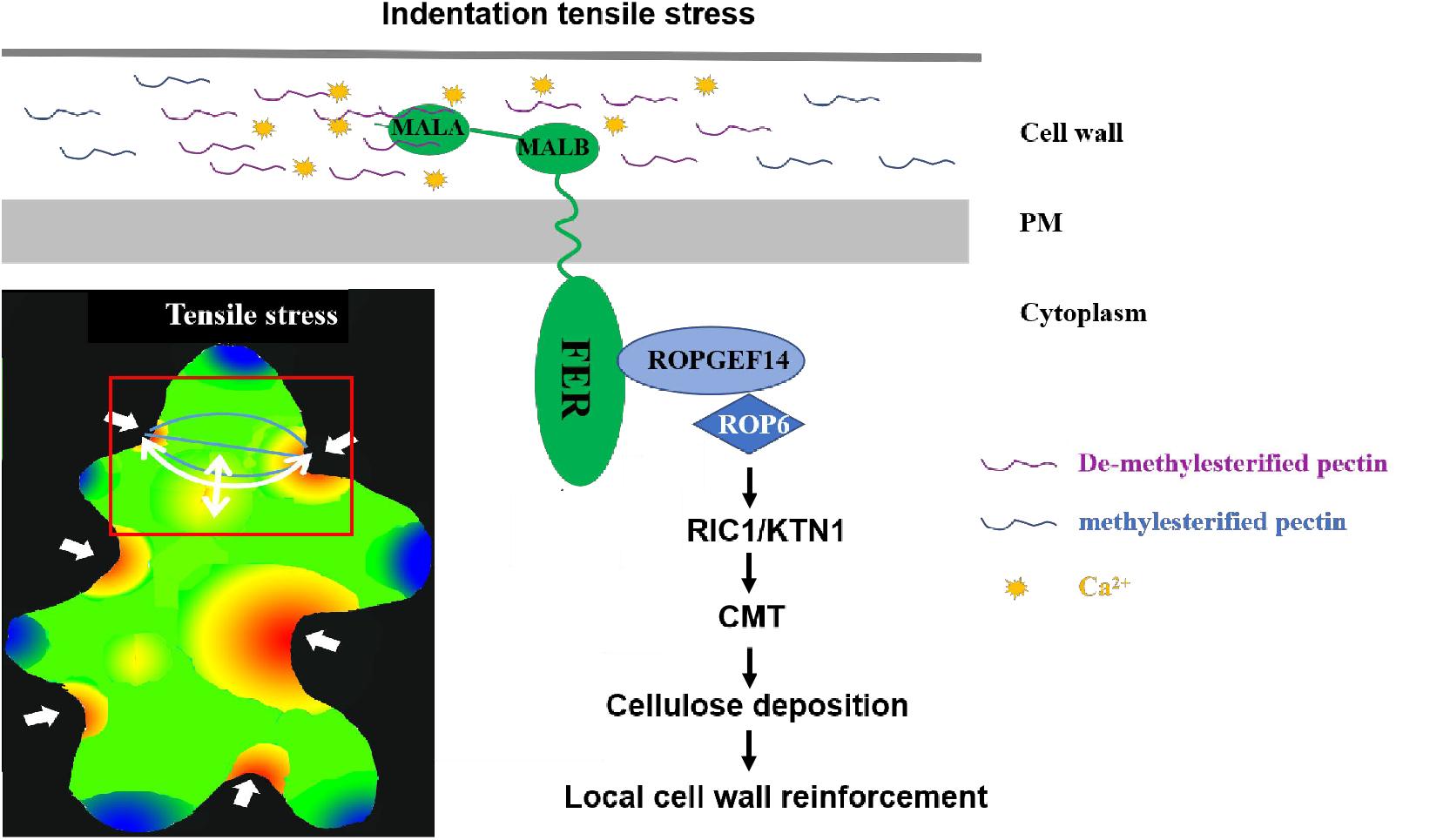
A working model for FER’s function in sensing/signaling mechanical stress in PC indentation region to strengthen local cell wall by activating the ROP6 signaling pathway. Indentation regions of PC experience highest mechanical stress [33]. By FER binding to highly demethylesterified pectin, which preferentially accumulates in the indentation region, FER may sense this mechanical stress and activates the ROP6 signaling pathway by interacting with ROPGEF14. Active ROP6 then promotes the reorganization of cortical microtubule (CMT), which guilds cellulose deposition to strengthen local cell wall, leading to increased stiffness of the cell wall. Left bottom arrowheads indicate regions of highest stress magnitude in neck regions, blue lines indicate radial arrangement of CMT and white double arrow lines shows the tensile stress.

Extracellular matrices (ECM) such as cell walls in plants and fungi have long been proposed to participate in mechanosensation, and this concept is well-established in animal systems [56, 57]. This concept involves the direct binding of an ECM component to the extracellular region of a transmembrane receptor, in such that the stiffness of ECM and changes in turgor pressure in the cell, could be mechanically monitored by the receptor *via* changes its conformation. However direct evidence for cell wall’s role in the mechanotransduction and the identity of its transmembrane receptor partner remained elusive. As in animals, the cytoskeleton is also a central feature of the plant cell’s response to mechanical stress [30], but how mechanical signals regulate the re-organization of the cytoskeleton is largely unknown. Therefore, our finding that the pectin-FER module is involved in mechanosensing and/or mechano-transduction and is directly linked to ROP6 signaling to the re-organization of cortical MTs is quite significant, as it provides important insights into the molecular mechanisms for mechanotransduction to regulate cell expansion and cell morphogenesis that relies on the mechanics of the cell wall in plants. Specifically, in PCs, the indenting regions are enriched in highly demethylesterified pectin making this region to experience mechanical stress differentially from lobing regions. Conceivably the pectin-FER senses the mechanical cue to promote cellulose microfibril deposition in the regions, re-enforcing them and enhancing the indentations (Figure 6). We propose that the pectin-FER system, by monitoring changes cell wall composition/modification and the resulting mechanical cues, feedback regulates the reconstruction of the cell wall, and consequently modulate cell shape formation, cell expansion, and cell wall integrity.

In addition to cell expansion and morphogenesis, FER has also been shown to maintain cell wall integrity under salt stress and in root hairs [45]. Several FER homologs in the CrRLK1L subfamily such as ANX1/2 and BUPS1/2 have been shown to maintain cell wall integrity in the rapidly growing cells such as pollen tubes. Interestingly BUPS1/CUP1 also binds pectin and participates in the sensing and/or signaling of mechanical stress to ROPs to control cell wall integrity of pollen tubes during their penetrative growth with the pistil [45, 58]. Hence the CrRLK1L subfamily of receptor kinases may have a widespread role in mechanosensation and/or mechano-transduction in plants.

## METHODS

### Plant Materials and Growth Conditions

The *fer-4* (GABI_GK106A06) and *gef14-2* (Salk_064617C) were obtained from the Arabidopsis Biological Resource Center (ABRC) at Ohio State University, Columbus, OH. The *PMEI1-OE* were obtained from Vincenzo Lionetti (Sapienza University of Rome, Italy). *GFP-TUBxrop6-1* and *GFP-TUBxric1-1* were generated in our lab as described in [29]. The *GFP-MAP4xfer-4, GFP-TUBxfer-4, GFP-ROP6xfer-4, GFP-ROP2xfer-4, GFP-MAP4xPMEI1OE, GFP-ROP6xPMEI1OE, GFP-ROP6xgef14-2* and *GFP-TUBxgef14-2* mutants were generated by genetic crosses and confirmed by genotyping or Western blotting. *Arabidopsis* plants were grown in soil (Sungro S16-281) in a growth room at 23°C, 40% relative humidity, and 75 μE m^-2^·s^-1^ light with a 12-h photoperiod. To grow *Arabidopsis* seedlings, the seeds were surface sterilized with 50% (vol/vol) bleach for 10 min, and then placed on the plates with ½ MS medium containing 0.5% sucrose, and 1.0% agar at pH 5.7.

### Mechanical perturbations

Three types of mechanical perturbations (ablation, compression and Isoxaben treatment) were applied on pavement cells of *Arabidopsis* cotyledons. Single pavement cell from the same region of cotyledon was ablated by high-dosage UV light radiation (Confocal SP5) on the epidermis of cotyledons from seedlings. When the target cell was ablated, there would be a dark region with clear cell shape left. Ablation time depended on the stage of cotyledons. For the ablation of 1 DAG PCs, 10s were sufficient. For 3 DAG PCs, we applied 30-40s of UV irradiation. For 7 DAG PCs, the ablation time was increased to 60s. For compression, we first moved seedlings to a *petri dish* plate with absorbent paper (kept wet with liquid ½ MS) and placed a coverslip beside them. Then we applied compressive forces on the cotyledons by using a coverslip that was pressed on the surface of the cotyledons and kept in place using adhesive silicone applied on the margins. Then the samples were loaded for MT observation. To investigate the effects of compression on ROP activity, 3-weeks old leaves were collected and cultured in a plate with absorbent paper (kept wet with liquid½ MS). After 30 min, four glass slides were placed on one half of a leaf as compression treatment, and the other half was free of external pressure as control. At various times, the glass slides were removed, and the leaves were dissected into two halves and immediately frozen in liquid nitrogen for ROP6 activity assay. For Isoxaben treatment on cotyledons for MT observation, seedlings were cultured in solid ½ MS and then moved to a plate with absorbent paper (kept wet with liquid ½ MS) for 30 min. The seedlings were then loaded for MT imaging on cotyledon PCs (0 h). After imaging these seedlings were transferred to a *petri dish* plate with absorbent paper, which was kept wet with liquid ½ MS containing Isoxaben. After 4 h or 6 h, the same cotyledons were loaded for MT imaging on the same regions imaged at 0 hr. For the analysis of Isoxaben treatment on ROP activities, seedlings were first cultured to 7 DAG in half solid ½ MS (0.4% agar), and then transferred to regular liquid ½ MS. After 30 min incubation, the seedlings were transferred to liquid ½ MS containing different concentrations of Isoxaben. Finally, the Isoxaben-treated seedlings were collected for ROP activity measurements.

### Quantitative analysis of cortical microtubule orientation

CMT of cotyledon PCs expressing MAP4-GFP were imaged using a Leica SP5 Laser Scanning Confocal. FibrilTool, an ImageJ plug-in to quantify fibrillar structures, was used to analyze the average anisotropy of MT [52]. The anisotropy score “0” indicates no order (isotropic arrays) and “1” indicates perfectly ordered (anisotropic arrays). For quantitative analysis of local cortical microtubule orientation in Figures 1C and 1D, only the indentation regions were selected with the Polygon tool for the analysis. Each data set was from the measurement of at least 20 cells collected from 3 different cotyledons from 3 individual seedlings. To examine the effect of mechanical stress on MT rearrangements, anisotropy of entire pavement cell before and after treatment was measured. For ablation treatment, two layers of PCs surrounding the ablated cell were used for anisotropy analysis. For compression and Isoxaben treatments, PCs at the same region of cotyledon (mid-region) were selected for anisotropy analysis.

### Measurement of ROP6 activity

Western blotting-based ROP activity measurements were performed as described in [24]. Total ROP6 and activated ROP6 proteins that were pulled down by MBP-RIC1 (GTP-bound ROP6) were detected by Western blotting analysis using anti-ROP6 or anti-GFP and horseradish peroxidase-conjugated rabbit antibodies. For ROP6 activity assays in 7 DAG seedlings, 0.2 g samples were used for each assay. For 3-week-old leaves, 0.4-0.5 g samples were collected for ROP activity measurements. Image J was used to quantify the mounts of ROPs.

## Acknowledgements

We thank Olivier Hamant and members of the Yang laboratory for stimulating discussion and critical comments on this work. We thank Vincenzo Lionetti (Sapienza University of Rome, Italy) for providing PMEI1OE seeds. We appreciate microscopy assistance from the Institute of Integrative Genome Biology Microscopy Core Faculty (UC Riverside, US). This work is in part supported by a grant from National Natural Science Foundation of China to W.T. (No. 3150080116).

## Supplemental Information

**Fig. S1.**
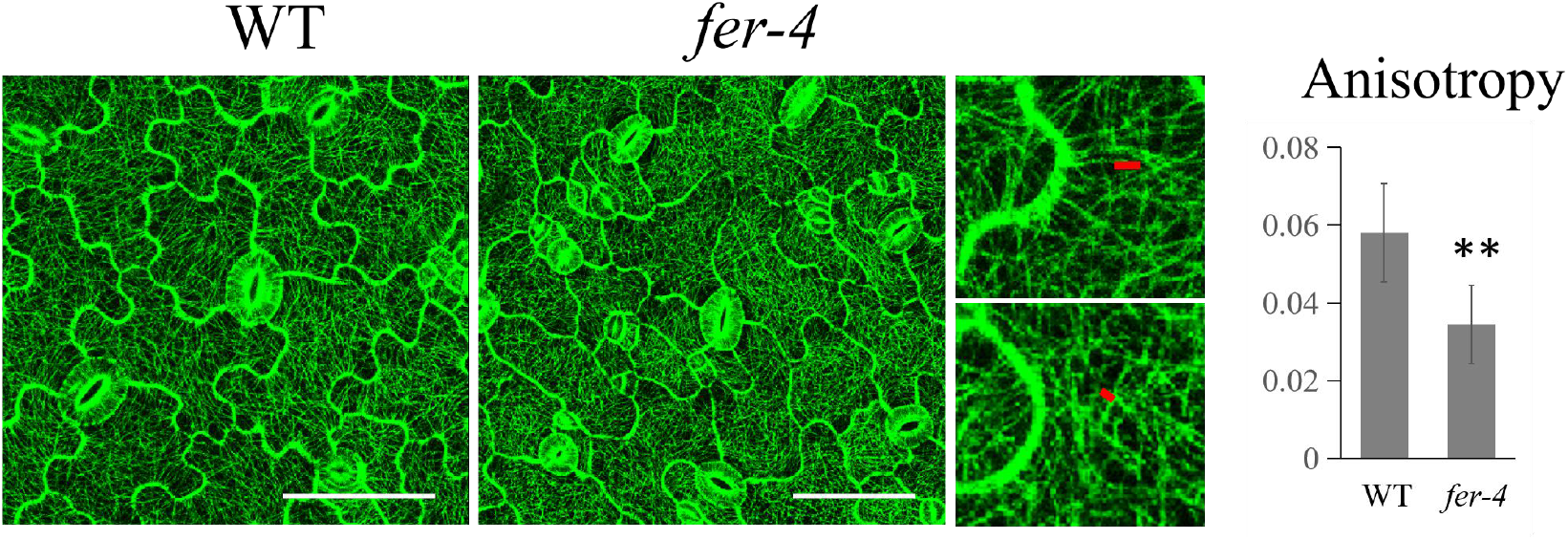
The radial distribution of cortical MT (microtubules) was greatly reduced in indentations of *fer-4* PCs (2 DAG). *Right* showed PC cortical MT of wild-type (GFP-MAP4) and the *fer-4* mutant (*fer-4xGFP-MAP4*) grown in ½ MS with 1.0 % agar. Bar, 50 μm. *Left* was the magnified PC indentation regions. The degree of cortical MT anisotropy was quantified. Red lines were generated by FibrilTool, an Image J plug-in to quantify fibrillar structures in raw microscopy images. the angle of red line represents the average orientation of the array and the length of red line is proportional to the array anisotropy. Data are mean degrees from >20 independent cells ±SE. \*\**p*<0.01

**Fig. S2.**
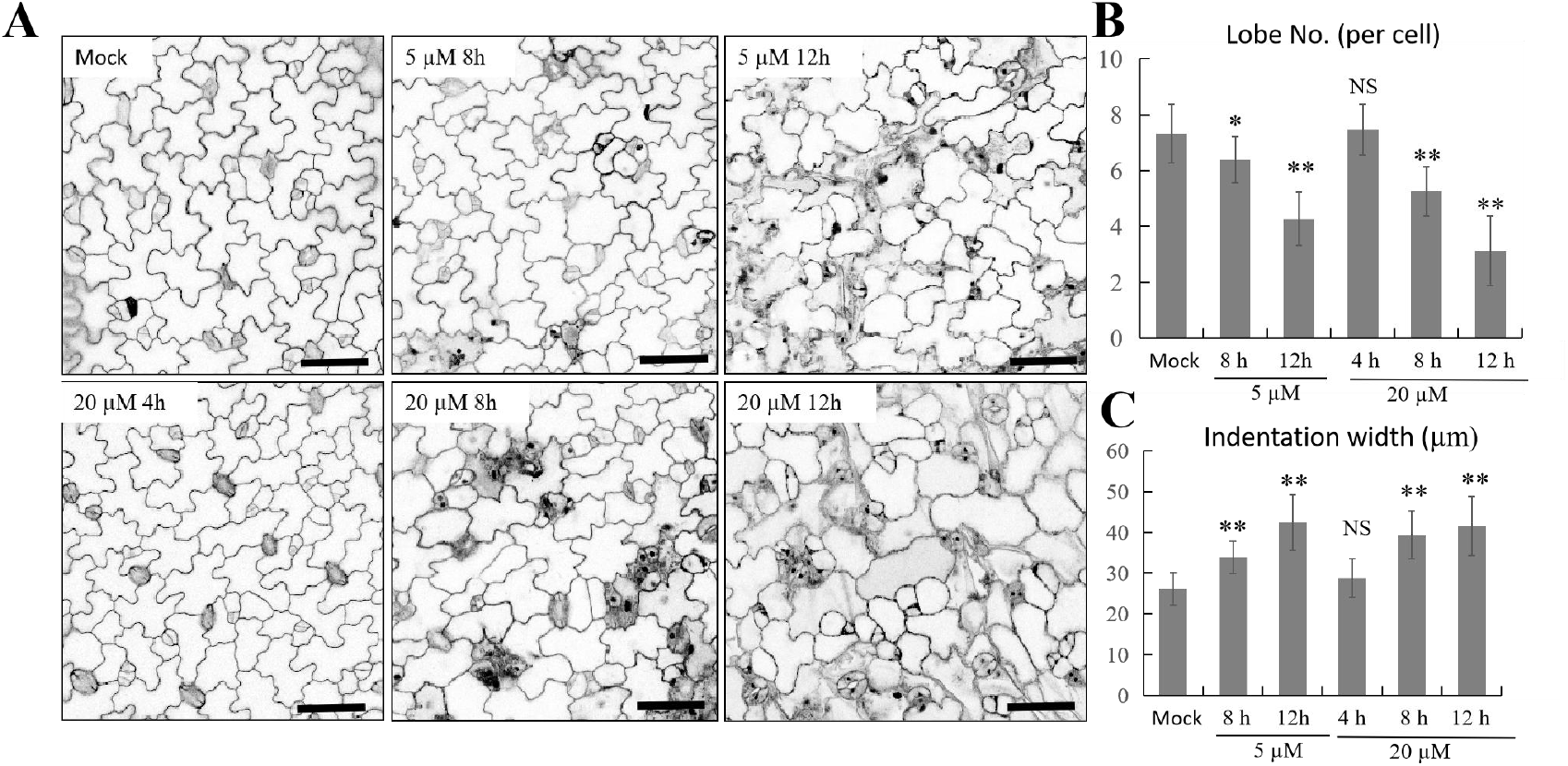
Isoxaben Treatment induces pavement cell shape change. (A) PC phenotypes of wild-type (Col-0) treated with different concentrations (mock, 5 μM, 20 μM of Isoxaben in different times (0 h, 4 h, 8 h, 12 h). Scale bar, 50 μm. (B) PC lobe number quantification result showed >5 μM Isoxaben significantly inhibit lobe formation. Data are represented as mean ± SE. \**p*<0.05; \*\**p*<0.01; Student’s *t*-test (C) PC indentation width quantification result showed >5 μM Isoxaben significantly inhibit neck formation. Data are represented as mean ± SE. \*\**p*<0.01; Student’s *t*-test

**Fig. S3.**
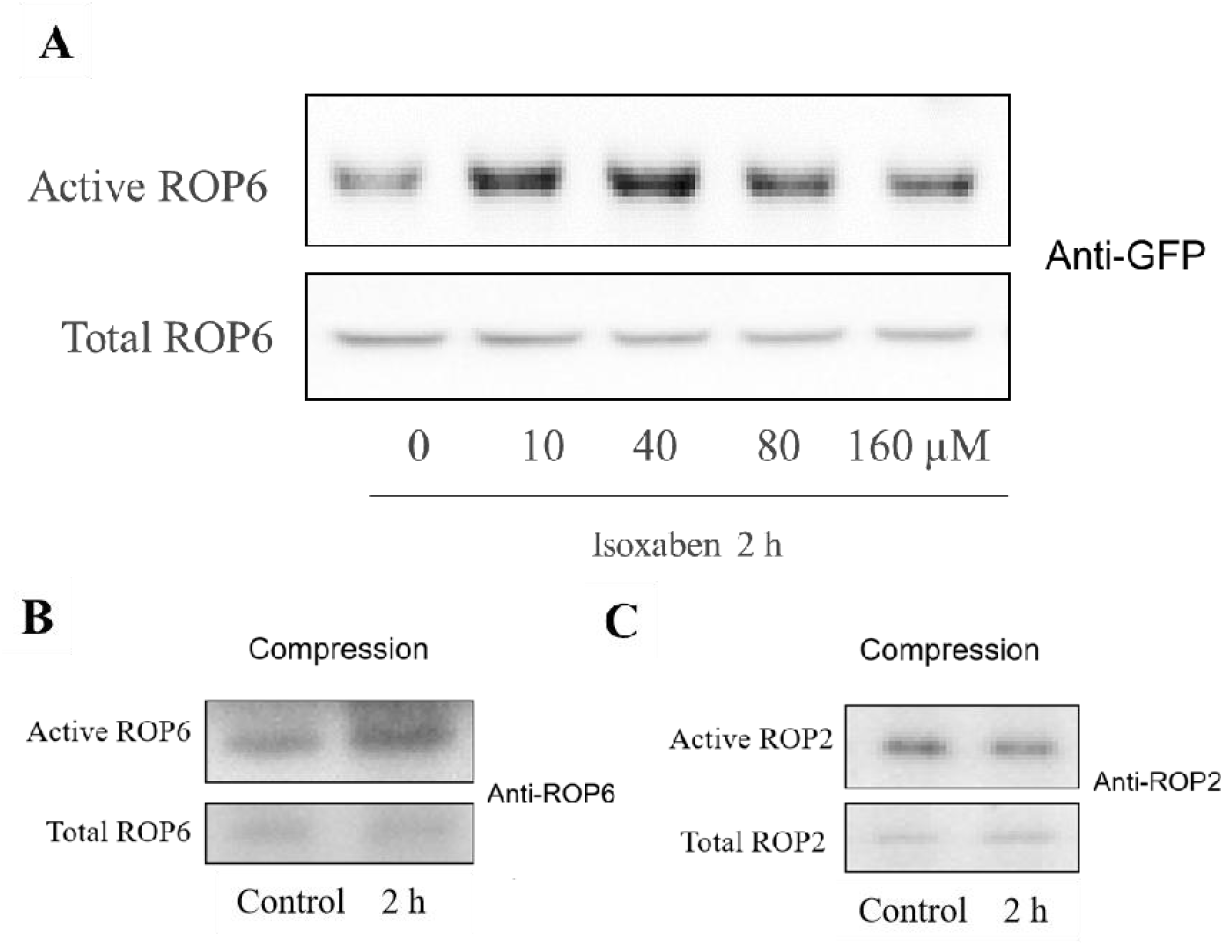
ROP6 can be activated by different concentrations of Isoxaben and compression. (A) ROP6 activity in 7 DAG seedlings of GFP-ROP6 with different concentrations of isoxaben. Seedlings of GFP-ROP6 were incubated in 1/2MS with 10, 20, 40, 80, 160 μM Isoxaben. The amount of active ROP6 increased most in 10, 40 μM Isoxaben treated seedlings comparing to 80 and 160 μM Isoxaben treatment (B) ROP6 is activated by compression. 3 weeks leaves of Col-0 were used for 2 hours compression treatment. Active and total ROP6 were detected by western blots using Anti-ROP6. (C) 2 hours compression have no effects on ROP2 activation. 3 weeks leaves of Col-0 were used for 2 hours compression treatment. Active and total ROP2 were detected by western blots using Anti-ROP2.

**Fig S4.**
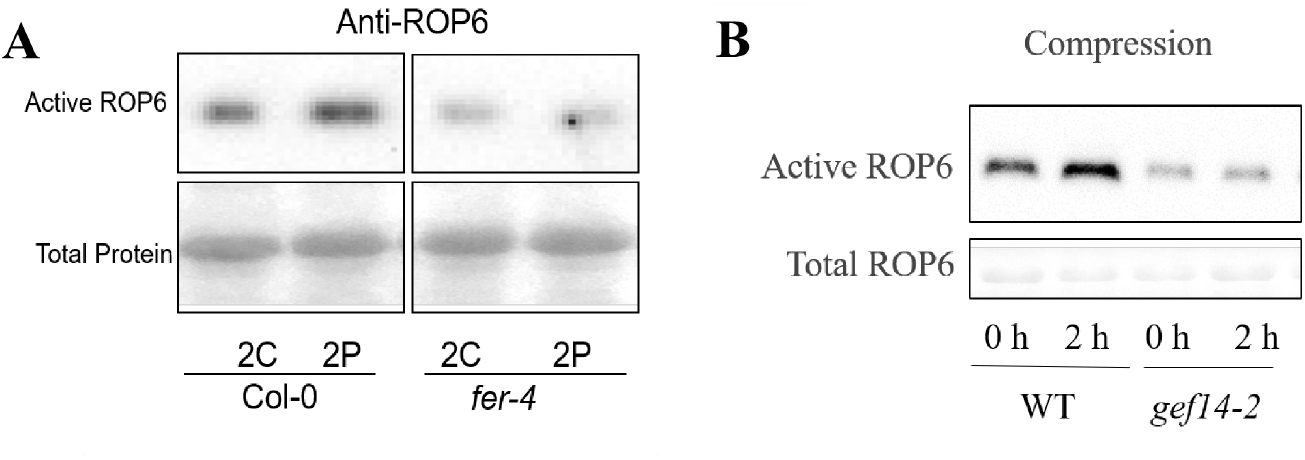
ROP6 can be obviously activated by compression in Col-0 but not *fer-4* and *gef14-2*. (A) Compression induced ROP6 activation is lost in *fer-4*. C, control; P, pressure (B) ROPGEF14 is required for compression induced ROP6 activation.

**Fig. S5.**
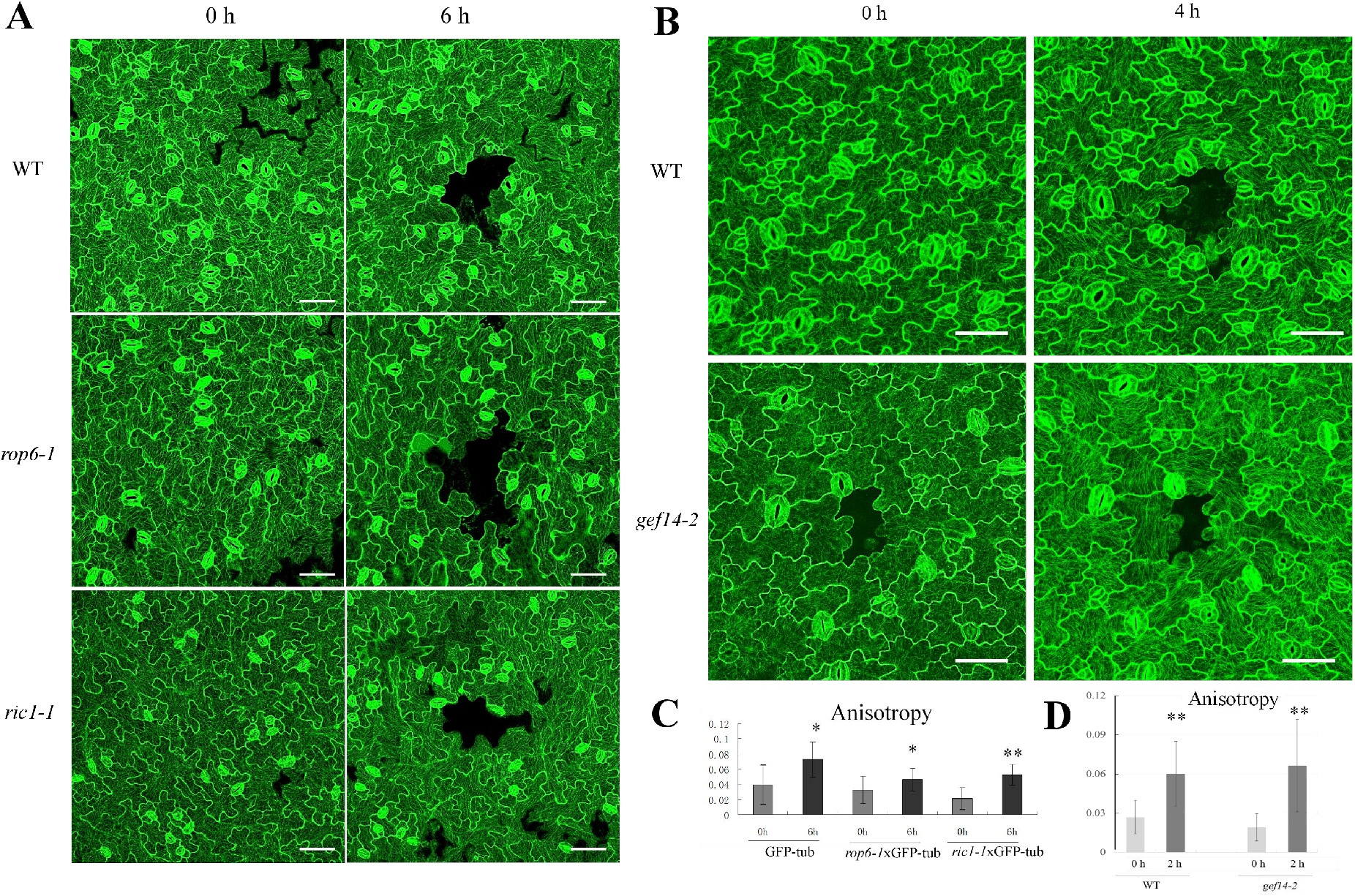
Early stage of PCs (3 DAG) of *rop6-1, ric1-1* and *gef14-2* showed normal sensitivity to ablation in terms of MT rearrangement. (A) MT rearrangements after single cell ablation in WT, *rop6-1* and *ric1-1* PCs. 7 DAG cotyledon grown in ½ MS with normal concentration of agar (1.0%) were observed in Confocal SP5. Red line represents the anisotropy of MT. Two layers of PCs around ablation point were quantified. Bar, 50 μm. (B) MT rearrangements after single cell ablation in WT and *gef14-2* PCs. 3 DAG cotyledon grown in ½ MS with normal concentration of agar (1.0%) were observed in Confocal SP5. Red line represents the anisotropy of MT. Two layers of PCs around ablation point were quantified. Bar, 50 μm. (C) Anisotropy value of MT in A. Significantly increased anisotropy means there is obvious MT rearrangement in WT PC after ablation. In normal concentration of agar, both *rop6-1* and *ric1-1* showed no significant differences comparing to WT in terms of anisotropy change. Data are mean degrees from >40 independent cells ±SE. *, p<0.05; **, p<0.01; NS, no significance (D) Anisotropy value of MT in B. *gef14-2* showed no significant differences comparing to WT in terms of anisotropy change. Data are mean degrees from >40 independent cells ±SE. **, p<0.01

